# Evaluation of the immunogenicity of prime-boost vaccination with the replication-deficient viral vectored COVID-19 vaccine candidate ChAdOx1 nCoV-19

**DOI:** 10.1101/2020.06.20.159715

**Authors:** Simon P. Graham, Rebecca K. McLean, Alexandra J. Spencer, Sandra Belij-Rammerstorfer, Daniel Wright, Marta Ulaszewska, Jane C. Edwards, Jack W. P. Hayes, Veronica Martini, Nazia Thakur, Carina Conceicao, Isabelle Dietrich, Holly Shelton, Ryan Waters, Anna Ludi, Ginette Wilsden, Clare Browning, Dagmara Bialy, Sushant Bhat, Phoebe Stevenson-Leggett, Philippa Hollinghurst, Ciaran Gilbride, David Pulido, Katy Moffat, Hannah Sharpe, Elizabeth Allen, Valerie Mioulet, Chris Chiu, Joseph Newman, Amin S. Asfor, Alison Burman, Sylvia Crossley, Jiandong Huo, Raymond J. Owens, Miles Carroll, John A. Hammond, Elma Tchilian, Dalan Bailey, Bryan Charleston, Sarah C. Gilbert, Tobias J. Tuthill, Teresa Lambe

## Abstract

Clinical development of the COVID-19 vaccine candidate ChAdOx1 nCoV-19, a replication-deficient simian adenoviral vector expressing the full-length SARS-CoV-2 spike (S) protein was initiated in April 2020 following non-human primate studies using a single immunisation. Here, we compared the immunogenicity of one or two doses of ChAdOx1 nCoV-19 in both mice and pigs. Whilst a single dose induced antigen-specific antibody and T cells responses, a booster immunisation enhanced antibody responses, particularly in pigs, with a significant increase in SARS-CoV-2 neutralising titres.

## Introduction

As SARS-CoV-2 began to spread around the world at the beginning of 2020 several vaccine platform technologies were employed to generate candidate vaccines. Several use replication-deficient adenoviral (Ad) vector technology and express the SARS-CoV-2 spike (S) protein. The first phase I clinical study of an Ad5-vectored vaccine has been reported^1^, ChAdOx1 nCoV-19 (AZD1222) phase I trials (NCT04324606) began in April 2020 with phase II and III trials (NCT04400838) started soon thereafter, and an Ad26-vectored vaccine is expected to enter phase I shortly. Typically, only one dose of Ad-vectored vaccines has been administered in early preclinical challenge studies or clinical studies against emerging or outbreak pathogens^2-5^. Rhesus macaques immunised with a single dose of ChAdOx1 nCoV-19 were protected against pneumonia but there was no impact on nasal virus titers after high dose challenge to both the upper and lower respiratory tract^6^. To increase antibody titres and longevity of immune responses, a booster vaccination may be administered. Homologous prime-boost immunisation resulted in higher antibody titres including neutralising antibodies and a trend towards a lower clinical score in a MERS-CoV challenge study^7^. Here, we set out to test the immunogenicity of either one or two doses of ChAdOx1 nCoV-19 in mice and pigs, to further inform clinical development.

## Results

‘Prime-boost’ vaccinated inbred (BALB/c) and outbred (CD1) mice were immunised on 0 and 28 days post-vaccination (dpv), whereas, ‘prime-only’ mice received a single dose of ChAdOx1 nCoV-19 on day 28. Spleens and serum were harvested from all mice on day 49 (3 weeks after boost or prime vaccination). Analysis of SARS-CoV-2 S protein-specific murine splenocyte responses by IFN-γ ELISpot assay showed no statistically significant difference between the prime-only and prime-boost vaccination regimens, in either strain of mouse (Figure 1A). Intracellular cytokine staining (ICS) of splenocytes (Figure 1B) showed, in both mouse strains, that the response was principally driven by CD8^+^ T cells. The predominant cytokine response of both CD8^+^ and CD4^+^ T cells was expression of IFN-γ and TNF-α, with negligible frequencies of IL-4^+^ and IL-10^+^ cells, consistent with previous data suggesting adenoviral vaccination does not induce a dominant Th2 response^8,9^. There were no signficant differences in CD4^+^ and CD8^+^ T cell cytokine responses between prime-only and prime-boost mice. Prime-only and prime-boost pigs were immunised on 0 dpv and prime-boost pigs received a second immunisation on 28 dpv. Blood samples were collected weekly until 42 dpv to analyse immune responses. IFN-γ ELISpot analysis of porcine peripheral blood mononuclear cells (PBMC) showed responses on 42 dpv (2 weeks after boost) that were significantly greater in the prime-boost pigs compared to prime-only animals (*p* < 0.05; Figure 1C). The prime-boost 42 dpv responses were greater than responses observed in either group on 14 dpv, but inter-animal variation meant this did not achieve statistical significance. ICS analysis of porcine T cell reponses showed a dominance of Th1-type cytokines (similar to the murine response) but with a higher frequency of S-specific CD4^+^ T cells compared to CD8^+^ T cells (Figure 1D). However, CD4^+^ and CD8^+^ T cell cytokine responses did not differ significantly between vaccine groups or timepoints (14 vs. 42 dpv).

**Figure 1:**
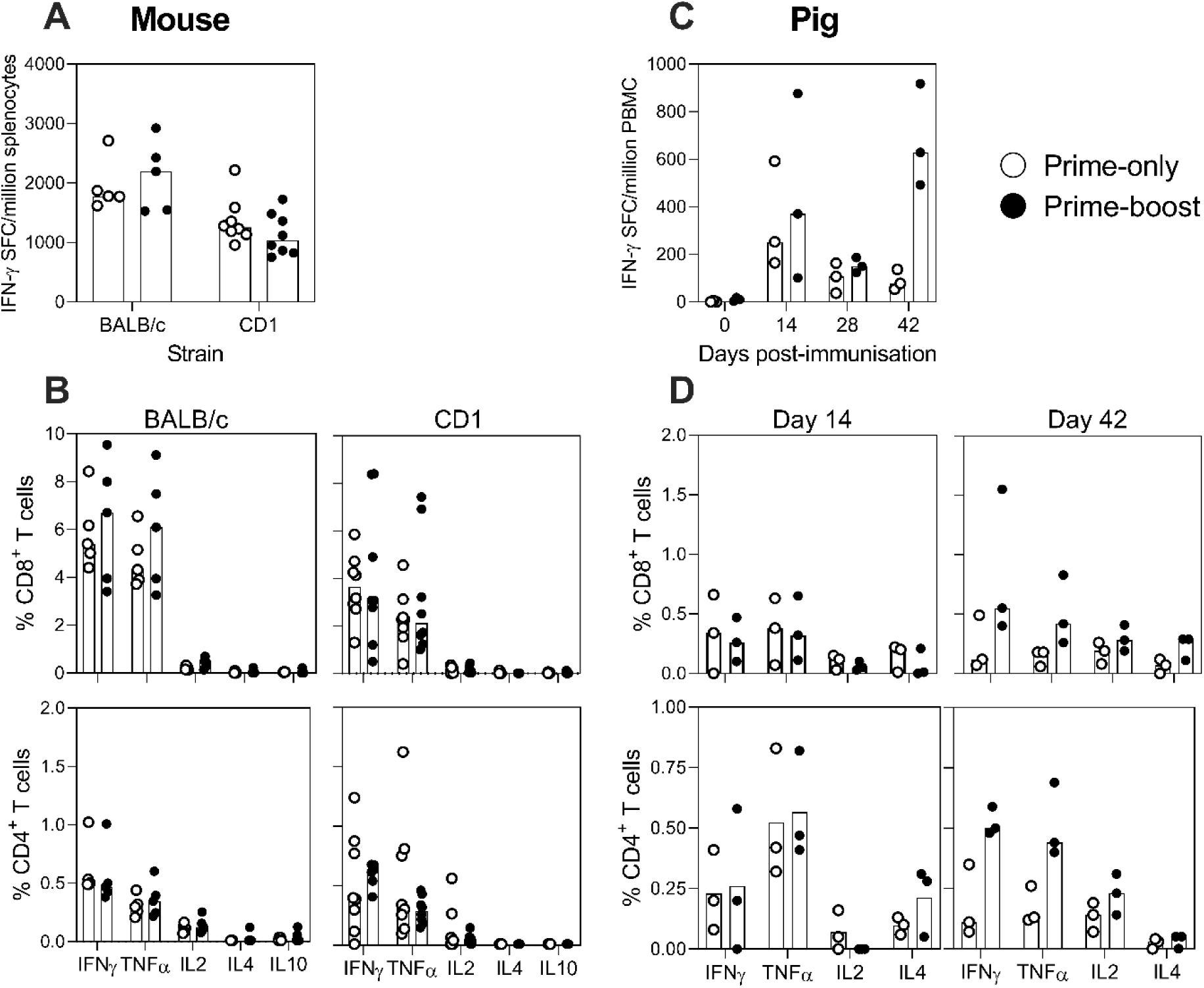
SARS-CoV-2 S-specific T cell responses following ChAdOx1 nCoV-19 prime-only and prime-boost vaccination regimens in mice and pigs. Inbred BALB/c (n=5) and outbred CD1 (n=8) were immunised on day 0 and 28 with ChAdOx1 nCoV19 (Prime-boost) or ChAdOx1 nCoV19 on day 28 (Prime-only); pigs (n=3) were immunised with ChAdOx1 nCoV-19 on days 0 and 28 (Prime-boost), or only on day 0 (Prime-only). To analyse SARS-CoV-2 S-specific T cell responses, all mice were sacrificed on day 49 for isolation of splenocytes and pigs were blood sampled longitudinally to isolate PBMC. Following stimulation with SARS-CoV-2 S-peptides, responses of murine splenocytes (**A**) and porcine PBMC (**C**) were assessed by IFN-γ ELISpot assays. Using flow cytometry, CD4^+^ and CD8^+^ T cell responses were characterised by assessing expression of IFN-γ, TNF-α, IL-2, IL-4 and IL-10 (mice; **B**) and IFN-γ, TNF-α, IL-2 and IL-4 (pigs; **D**). Each data point represents an individual mouse/pig with bars denoting the median response per group/timepoint.

SARS-CoV-2 S protein-specific antibody titres in serum were determined by ELISA using recombinant soluble trimeric S (FL-S) and receptor binding domain (RBD) proteins. A significant increase in FL-S binding antibody titres was observed in prime-boost BALB/c mice compared to their prime-only counterparts (*p* < 0.01), however, the difference between vaccine groups for CD1 mice was not significant (Figure 2A). Antibody responses were evaluated longitudinally in pig sera by FL-S and RBD ELISA. Compared to pre-vaccination sera, significant FL-S specific antibody titres were detected in both prime-only and prime-boost groups from 21 and 14 dpv, respectively (*p* < 0.01; Figure 2B). FL-S antibody titres did not differ signifcantly between groups until after the boost, when titres in the prime-boost pigs became significantly greater with an average increase in titres of > 1 log_10_ (*p* < 0.0001). RBD-specific antibody titres showed a similar profile with significant titres in both groups from 14 dpv (*p* < 0.05) and a further significant increase in the prime-boost pigs from 35 dpv onwards which was greater than the prime-only pigs (*p* < 0.0001; Figure 2C). SARS-CoV-2 neutralising antibody responses were assessed using a virus neutralisation test (VNT; Figure 2D) and pseudovirus-based neutralisation test (pVNT; Figure 2E). After the prime immunisation, SARS-CoV-2 neutralising antibody titres were detected by VNT in 14 and 28 dpv sera from 2/3 prime-boost and 2/3 prime-only pigs. Two weeks after the boost (42 dpv), neutralising antibody titres were detected and had increased in all prime-boost pigs, which were significantly greater than the earlier timepoints and the titres measured in the prime-only group (*p* < 0.01). In agreement with this analysis, serum assayed for neutralising antibodies using the pVNT revealed that antibody titres in 42 dpv prime-boost pig sera were significantly greater than earlier timepoints and the prime-only group (*p* < 0.001). Statistical analysis showed a highly significant correlation between pVNT and VNT titres (Spearman’s rank correlation *r* = 0.86; *p* < 0.0001).

**Figure 2:**
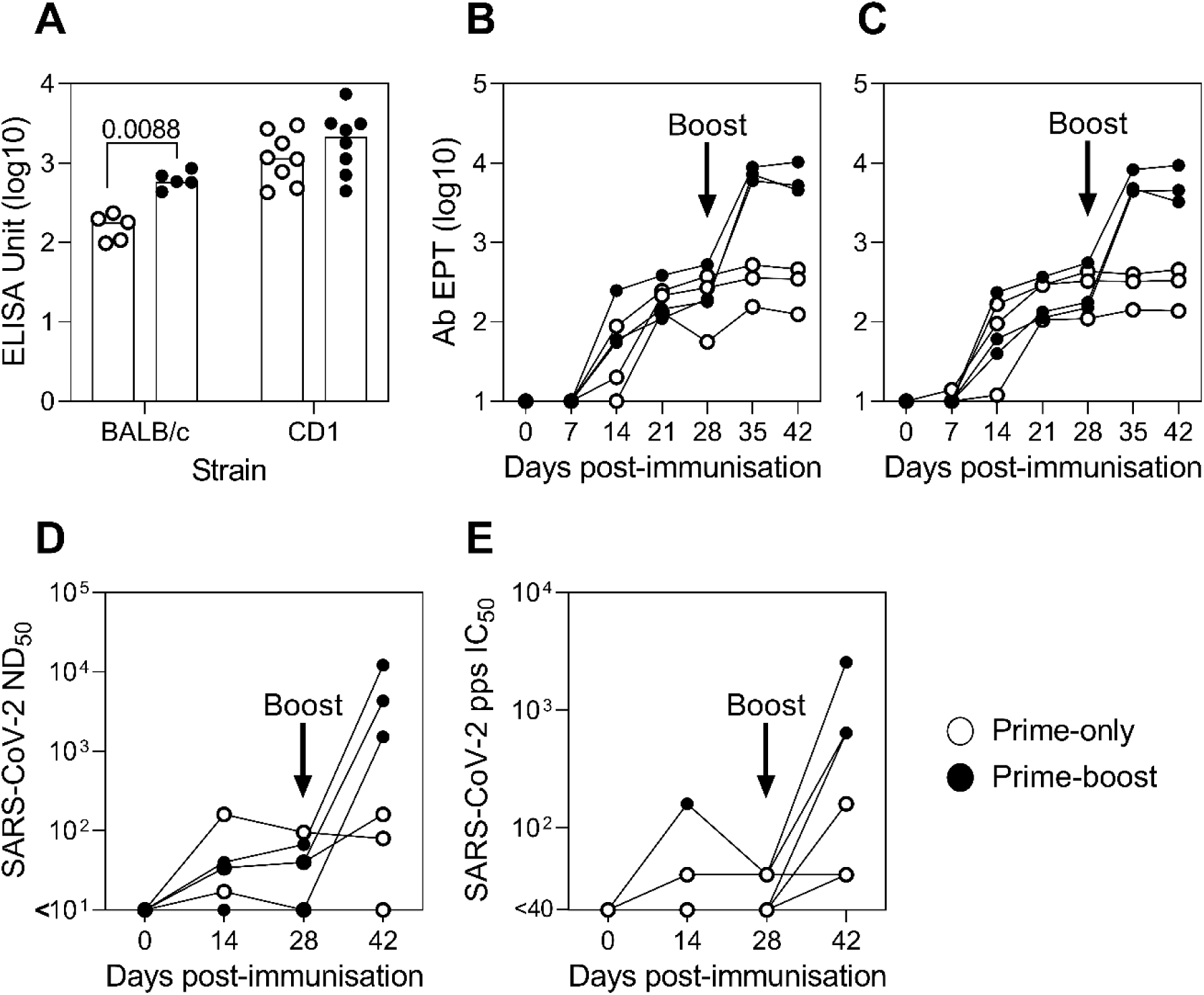
SARS-CoV-2 S protein-specific antibody responses following ChAdOx1 nCoV-19 prime-only and prime-boost vaccination regimens in mice and pigs. Inbred BALB/c (n=5) and outbred CD1 (n=8) were immunised on day 0 and 28 with ChAdOx1 nCoV19 (Prime-boost) or ChAdOx1 nCoV19 on day 28 (Prime-only), whereas, pigs were immunised with ChAdOx1 nCoV-19 on days 0 and 28 (Prime-boost), or only on day 0 (Prime-only). To analyse SARS-CoV-2 S protein-specific antibodies in serum, all mice were sacrificed on day 49 and pigs were blood sampled weekly until day 42. Antibody units or end-point titres (EPT) were assessed by ELISA using recombinant SARS-CoV-2 FL-S for both mice (**A**) and pigs (**B**), and recombinant S protein RBD for pigs (**C**). SARS-CoV-2 neutralising antibody titres in pig sera were determined by VNT, expressed as the reciprocal of the serum dilution that neutralised virus infectivity in 50% of the wells (ND_50_; **D**), and pVNT, expressed as reciprocal serum dilution to inhibit pseudovirus entry by 50% (IC_50_; **E**). Each data point represents an individual mouse/pig sera with bars denoting the median titre per group.

## Discussion

In this study, we utilised both a small and a large animal model to evaluate the immunogenicity of either one or two doses of a COVID-19 vaccine candidate, ChAdOx1 nCoV-19 (now known as AZD1222). Small animal models have variable success in predicting vaccine efficacy in larger animals but are an important stepping stone to facilitate prioritisation of vaccine targets. In contrast, larger animal models, such as the pig and non-human primates, have been shown to more accurately predict vaccine outcome in humans^10-12^. The mouse data generated in this study suggested that the immunogenicity profile was at the upper end of a dose response curve, which may have saturated the immune response and largely obscured our ability to determine differences between prime-only or prime-boost regimens. We have developed the pig as a model for generating and understanding immune responses to vaccination against human influenza^13-15^ and Nipah virus^16,17^.The inherent heterogeneity of an outbred large animal model is more representative of immune responses in humans. Extensive development of reagents to study immune responses in pigs in recent years has extended the usefulness and applicability of the pig as a model to study infectious disease. These data demonstrate the utility of the pig as a model for further evaluation of the immunogenicity of ChAdOx1 nCoV-19 and other COVID-19 vaccines.

We show here that T cell responses are higher in pigs that received a prime-boost vaccination when compared to prime only at day 42, whilst comparing responses 14 days after last immunisation demonstrates the prime-boost regimen trended toward a higher response. In addition, ChAdOx1 nCoV-19 immunisation induced robust Th1-like CD4^+^ and CD8^+^ T cell responses in both pigs and mice. This has important implications for COVID-19 vaccine development as virus-specific T cells are thought to play an important role in SARS-CoV-2 infection ^18-22^. While no correlate of protection has been defined for COVID-19, recent publications suggest that neutralising antibody titres may be correlated with protection in animal challenge models^23,24^. A single dose of ChAdOx1 nCoV-19 induces antibody responses, but we demonstrate here that antibody responses are significantly enhanced after homologous boost in one mouse strain and to a greater extent in pigs. However, it is likely that a combination of neutralising antibodies and antigen-specific T cells would act in synergy to prevent and control infection, as we have recently shown in the context of influenza vaccination^13,25^.

Whilst human immunogenicity and clinical read-outs are a critically meaningful endpoint, studies in small animals and pigs will help prioritise candidates to be tested in humans. Further clinical studies are needed to assess immunogenicity after prime-boost vaccination and the impact on clinical efficacy and durability of the immune response.

## Methods

### Ethics statement

Mouse and pig studies were performed in accordance with the UK Animals (Scientific Procedures) Act 1986 and with approval from the relevant local Animal Welfare and Ethical Review Body (Mice – Project License P9808B4F1, and pigs – Project License PP1804248). The principles of the 3R’s were applied for the duration of the study to ensure animal welfare was not unnecessarily compromised.

### Cells and viruses

Vero E6 cells were grown in DMEM containing sodium pyruvate and L-glutamine (Sigma-Aldrich, Poole, UK), 10% FBS (Gibco, Thermo Fisher, Loughborough, UK), 0.2% penicillin/streptomycin (10,000 U/mL; Gibco) (maintenance media) at 37 °C and 5% CO_2_. SARS-CoV-2 isolate England-2 stocks were grown in Vero E6 cells using a multiplicity of infection (MOI) of 0.0001 for 3 days at 37 °C in propagation media (maintenance media containing 2% FBS). SARS-CoV-2 stocks were titrated on Vero E6 cells using MEM (Gibco), 2% FCS (Labtech, Heathfield, UK), 0.8% Avicel (FMC BioPolymer, Girvan, UK) as overlay. Plaque assays were fixed using formaldehyde (VWR, Leighton Buzzard, UK) and stained using 0.1% Toluidine Blue (Sigma-Aldrich). All work with live SARS-CoV-2 virus was performed in ACDP HG3 laboratories by trained personnel. The propagation, purification and assessment of ChAdOx1 nCoV-19 titres were as described previously^7^.

### Recombinant SARS-CoV-2 proteins and synthetic peptides

A synthetic DNA, encoding the spike (S) protein receptor binding domain (RBD; amino acids 330-532) of SARS-CoV-2 (GenBank MN908947), codon optimised for expression in mammalian cells (IDT Technology) was inserted into the vector pOPINTTGneo incorporating a C-terminal His6 tag. Recombinant RBD was transiently expressed in Expi293™ (Thermo Fisher Scientific, UK) and protein purified from culture supernatants by immobilised metal affinity followed by a gel filtration in phosphate-buffered saline (PBS) pH 7.4 buffer. A soluble trimeric S (FL-S) protein construct encoding residues 1-1213 with two sets of mutations that stabilise the protein in a pre-fusion conformation (removal of a furin cleavage site and the introduction of two proline residues; K983P, V984P) was expressed as described^26^. The endogenous viral signal peptide was retained at the N-terminus (residues 1-14), a C-terminal T4-foldon domain incorporated to promote association of monomers into trimers to reflect the native transmembrane viral protein, and a C-terminal His6 tag included for nickel-based affinity purification. Similar to recombinant RBD, FL-S was transiently expressed in Expi293™ (Thermo Fisher Scientific) and protein purified from culture supernatants by immobilised metal affinity followed by gel filtration in Tris-buffered saline (TBS) pH 7.4 buffer.

For analysis of T cell responses in pigs, overlapping 16mer peptides offset by 4 residues based on the predicted amino acid sequence of the entire S protein from SARS-CoV-2 Wuhan-Hu-1 isolate (NCBI Reference Sequence: NC_045512.2) were designed and synthesised (Mimotopes, Melbourne, Australia) and reconstituted in sterile 40% acetonitrile (Sigma-Aldrich) at a concentration of 3 mg/mL. Three pools of synthetic peptides representing residues 1-331 (Pool 1), 332-748 (Pool 2) and 749-1273 (Pool 3) were prepared for use to stimulate T cells in IFN-γ ELISpot and intracellular cytokine staining (ICS) assays. For analysis of T cell responses in mice, overlapping 15mer peptides offset by 11 residues were designed and synthesised (Mimotopes) and reconstituted in sterile 100% DMSO (Sigma-Aldrich) at a concentration of 100 mg/mL. Two peptide pools spanning S1 region (Pool 1: 1 to 77 and 317-321, Pool 2:78-167) and 2 peptide pools spanning S2 region (Pool 3:166 to 241, Pool 4:242 to 316) were used for stimulating splenocytes for IFN-γ ELISpot analysis, and single pools of S1 (Pool 1 and Pool 2) and S2 (Pool 3 and Pool 4) were used to stimulate splenocytes for ICS.

### Immunogenicity trials

#### Mice

Inbred female BALB/cOlaHsd (BALB/c) (Envigo) and outbred Crl:CD1 (CD1) (Charles River) of at least 6 weeks of age were randomly allocated into ‘prime-only’ or ‘prime-boost’ vaccination groups (BALB/c n=5 and CD1 n=8). Prime-boost mice were immunised intramuscularly with 10^8^ infectious units (IU) (6.02×10^9^ virus particles; vp) ChAdOx1 nCoV-19 and boosted intramuscularly four weeks later with 1 × 10^8^ IU ChAdOx1 nCoV-19. Prime-only mice received a single dose of 10^8^ IU ChAdOx1 nCoV-19 at the same time prime-boost mice were boosted. Spleens and serum were harvested from all animals a further 3 weeks later.

#### Pigs

Six 8–10-week-old, weaned, female, Large White-Landrace-Hampshire cross-bred pigs from a commercial rearing unit were randomly allocated to two treatment groups (*n* = 3): ‘Prime-only’ and ‘Prime-boost’. Both groups were immunised on day 0 with 1 × 10^9^ IU (5.12 × 10^10^ vp) ChAdOx1 nCoV-19 in 1 mL PBS by intramuscular injection (brachiocephalic muscle). ‘Prime-boost’ pigs received an identical booster immunisation on day 28. Blood samples were taken from all pigs on a weekly basis at 0, 7, 14, 21, 28, 35 and 42 dpv by venepuncture of the external jugular vein: 8 mL/pig in BD SST vacutainer tubes (Fisher Scientific) for serum collection and 40 mL/pig in BD heparin vacutainer tubes (Fisher Scientific) for peripheral blood mononuclear cell (PBMC) isolation.

### Detection of SARS-CoV-S-Specific Antibodies by ELISA

#### Mice

Antibodies to SARS-CoV-2 FL-S protein were determined by performing a standardised ELISA on serum collected 3-weeks after prime or prime-boost vaccination. MaxiSorp plates (Nunc) were coated with 100 ng/well FL-S protein overnight at 4°C, prior to washing in PBS/Tween (0.05% v/v) and blocking with Blocker Casein in PBS (Thermo Fisher Scientific) for 1 hour at room temperature (RT). Standard positive serum (pool of mouse serum with high endpoint titre against FL-S protein), individual mouse serum samples, negative and an internal control (diluted in casein) were incubated for 2 hours at RT. Following washing, bound antibodies were detected by addition of alkaline phosphatase-conjugated goat anti-mouse IgG (Sigma-Aldrich), diluted 1/5000 in casein, for 1 hour at RT and detection of anti-mouse IgG by the addition of pNPP substrate (Sigma-Aldrich). An arbitrary number of ELISA units were assigned to the reference pool and OD values of each dilution were fitted to a 4-parameter logistic curve using SOFTmax PRO software. ELISA units were calculated for each sample using the OD values of the sample and the parameters of the standard curve.

#### Pigs

Serum was isolated by centrifugation of SST tubes at 1300 × *g* for 10 minutes at RT and stored at −80°C. SARS-CoV-2 RBD and FL-S specific antibodies in serum were assessed as detailed previously^26^ with the exception of the following two steps. The conjugated secondary antibody was replaced with goat anti-porcine IgG HRP (Abcam, Cambridge, UK) at 1/10,000 dilution in PBS with 0.1% Tween_20_ and 1% non-fat milk. In addition, after the last wash, a 100 µL of TMB (One Component Horse Radish Peroxidase Microwell Substrate, BioFX, Cambridge Bioscience, Cambridge, UK) was added to each well and the plates were incubated for 7 minutes at RT. A 100 µL of BioFX 450nmStop Reagent (Cambridge Bioscience) was then added to stop the reaction and microplates were read at 450nm. End-point antibody titres (mean of duplicates) were calculated as follows: the log_10_ OD was plotted against the log_10_ sample dilution and a regression analysis of the linear part of this curve allowed calculation of the endpoint titre with an OD of twice the average OD values of 0 dpv sera.

### Assessment of SARS-CoV-2 Neutralising Antibody Responses

The ability of pig sera to neutralise SARS-CoV-2 was evaluated using virus and pseudovirus neutralisation assays. For both assays, sera were first heat-inactivated (HI) by incubation at 56 °C for 2 hours.

Virus neutralization test (VNT): Starting at a 1 in 5 dilution, two-fold serial dilutions of sera were prepared in 96 well round-bottom plates using DMEM containing 1% FBS and 1% Antibiotic-Antimycotic (Gibco) (dilution media). 75 μL of diluted pig serum was mixed with 75 μL dilution media containing approximately 64 plaque-forming units (pfu) SARS-CoV-2 for 1 hour at 37 °C. Vero E6 cells were seeded in 96-well flat-bottom plates at a density of 1 × 10^5^ cells/mL in maintenance media one day prior to experimentation. Culture supernatants were replaced by 100 µL of DMEM containing 10% FCS and 1% Antibiotic-Antimycotic, before 100 µl of the virus-sera mixture was added to the Vero E6 cells and incubated for six days at 37 °C. Cytopathic effect (CPE) was investigated by bright-field microscopy. Cells were further fixed and stained as described above, and CPE scored. Each individual pig serum dilution was tested in quadruplet on the same plate and no sera/SARS-CoV-2 virus and no sera/no virus controls were run concurrently on each plate in quadruplet. Wells were scored for cytopathic effect and neutralisation titres expressed as the reciprocal of the serum dilution that completely blocked CPE in 50% of the wells (ND_50_). Researchers performing the VNTs were blinded to the identity of the samples.

Pseudovirus neutralisation test (pVNT): Lentiviral-based SARS-CoV-2 pseudoviruses were generated in HEK293T cells incubated at 37 °C, 5% CO_2_. Cells were seeded at a density of 7.5 *x* 10^5^ in 6 well dishes, before being transfected with plasmids as follows: 500 ng of SARS-CoV-2 spike, 600 ng p8.91 (encoding for HIV-1 gag-pol), 600 ng CSFLW (lentivirus backbone expressing a firefly luciferase reporter gene), in Opti-MEM (Gibco) along with 10 µL PEI (1 µg/mL) transfection reagent. A ‘no glycoprotein’ control was also set up using carrier DNA (pcDNA3.1) instead of the SARS-CoV-2 S expression plasmid. The following day, the transfection mix was replaced with 3 mL DMEM with 10% FBS (DMEM-10%) and incubated at 37 °C. At both 48 and 72 hours post transfection, supernatants containing pseudotyped SARS-CoV-2 (SARS-CoV-2 pps) were harvested, pooled and centrifuged at 1,300 x *g* for 10 minutes at 4 °C to remove cellular debris. Target HEK293T cells, previously transfected with 500 ng of a human ACE2 expression plasmid (Addgene, Cambridge, MA, USA) were seeded at a density of 2 × 10^4^ in 100 µL DMEM-10% in a white flat-bottomed 96-well plate one day prior to harvesting of SARS-CoV-2 pps. The following day, SARS-CoV-2 pps were titrated 10-fold on target cells, with the remainder stored at −80 °C. For pVNTs, pig sera were diluted 1:20 in serum-free media and 50 µL was added to a 96-well plate in quadruplicate and titrated 4-fold. A fixed titred volume of SARS-CoV-2 pps was added at a dilution equivalent to 10^6^ signal luciferase units in 50 µL DMEM-10% and incubated with sera for 1 hour at 37 °C, 5% CO_2_. Target cells expressing human ACE2 were then added at a density of 2 *x* 10^4^ in 100 µL and incubated at 37 °C, 5% CO_2_ for 72 hours. Firefly luciferase activity was then measured with BrightGlo luciferase reagent and a Glomax-Multi^+^ Detection System (Promega, Southampton, UK). Pseudovirus neutralization titres were expressed as the reciprocal of the serum dilution that inhibited luciferase expression by 50% (IC_50_).

### Assessment of SARS-CoV-2 Specific T cell Responses

#### Mice

Single cell suspension of mouse spleens were prepared by passing cells through 70 μm cell strainers and ACK lysis (Thermo Fisher) prior to resuspension in complete media (αMEM supplemented with 10% FCS, Pen-Step, L-Glut and 2-mercaptoethanol). For analysis of IFN-γ production by ELISpot assay, splenocytes were stimulated with S peptide pools at a final concentration of 2μg/ml on IPVH-membrane plates (Millipore) coated with 5μg/ml anti-mouse IFN-γ (clone AN18; Mabtech). After 18-20 hours of stimulation, IFN-γ spot forming cells (SFC) were detected by staining membranes with anti-mouse IFN-γ biotin mAb (1 µg/mL; clone R46A2, Mabtech) followed by streptavidin-alkaline phosphatase (1 µg/mL) and development with AP conjugate substrate kit (Bio-Rad). For analysis of intracellular cytokine production, cells were stimulated with 2 μg/mL S peptide pools, media or cell stimulation cocktail (containing PMA-Ionomycin, BioLegend), together with 1 μg/mL GolgiPlug (BD Biosciences) and 2 μL/mL CD107a-Alexa647 for 6 hours in a 96-well U-bottom plate, prior to placing at 4°C overnight. Following surface staining with CD4-BUV496, CD8-PerCP-Cy5.5, CD62L-BV711, CD127-BV650, CD44-APC-Cy7 and LIVE/DEAD Aqua (Thermo Fisher), cells were fixed with 10% neutral buffered formalin (containing 4% paraformaldehyde) and stained intracellularly with TNF-α-AF488, IL-2-PE-Cy7, IL-4-BV605, IL-10-PE and IFN-γ-e450 diluted in Perm-Wash buffer (BD Biosciences). Sample acquisition was performed on a Fortessa (BD) and data analysed in FlowJo v9 (TreeStar). An acquisition threshold was set at a minimum of 5000 events in the live CD3^+^ gate. Antigen-specific T cells were identified by gating on LIVE/DEAD negative, doublet negative (FSC-H vs FSC-A), size (FSC-H vs SSC), CD3^+^, CD4^+^ or CD8^+^ cells and cytokine positive. Total SARS-CoV-2 S specific cytokine responses are presented after subtraction of the background response detected in the media stimulated control spleen sample of each mouse, prior to summing together the frequency of S1 and S2 specific cells.

#### Pigs

PBMCs were isolated from heparinised blood by density gradient centrifugation and cryopreserved in cold 10% DMSO (Sigma-Aldrich) in HI FBS^17^. Resuscitated PBMC were suspended in RPMI 1640 medium, GlutaMAX supplement, HEPES (Gibco) supplemented with 10 % HI FBS (New Zealand origin, Life Science Production, Bedford, UK), 1% Penicillin-Streptomycin and 0.1% 2-mercaptoethanol (50 mM; Gibco) (cRPMI). To determine the frequency of SARS-CoV-2 S specific IFN-γ producing cells, an ELISpot assay was performed on PBMC from 0, 14, 28 and 42 dpv. Multiscreen 96-well plates (MAHAS4510; Millipore, Fisher Scientific) were pre-coated with 1 µg/mL anti-porcine IFN-γ mAb (clone P2G10, BD Biosciences) and incubated overnight at 4 °C. After washing and blocking with cRPMI, PBMCs were plated at 5 × 10^5^ cells/well in cRPMI in a volume of 50 µL/well. PBMCs were stimulated in triplicate wells with the SARS-CoV-2 S peptide pools at a final concentration of 1 µg/mL/peptide. cRPMI alone was used in triplicate wells as a negative control. After 18 hours incubation at 37 °C with 5% CO_2_, plates were developed as described previously^17^. The numbers of specific IFN-γ secreting cells were determined using an ImmunoSpot^®^ S6 Analyzer (Cellular Technology, Cleveland, USA). For each animal, the mean ‘cRPMI only’ data was subtracted from the S peptide pool 1, 2 and 3 data which were then summed and expressed as the medium-corrected number of antigen-specific IFN-γ secreting cells per 1 *×* 10^6^ PBMC. To assess intracellular cytokine expression PBMC from 14 and 42 dpv were suspended in cRPMI at a density of 2 × 10^7^ cells/mL and added to 50 µL/well to 96-well round bottom plates. PBMCs were stimulated in triplicate wells with the SARS-CoV-2 S peptide pools (1 µg/mL/peptide). Unstimulated cells in triplicate wells were used as a negative control. After 14 hours incubation at 37 °C, 5% CO_2_, cytokine secretion was blocked by addition 1:1,000 BD GolgiPlug (BD Biosciences) and cells were further incubated for 6 hours. PBMC were washed in PBS and surface labelled with Zombie NIR fixable viability stain (BioLegend), CD4-PerCP-Cy5.5 mAb (clone 74-12-4, BD Bioscience) and CD8β-FITC mAb (clone PPT23, Bio-Rad Antibodies). Following fixation (Fixation Buffer, BioLegend) and permeabilization (Permeabilization Wash Buffer, BioLegend), cells were stained with: IFN-γ-AF647 mAb (clone CC302, Bio-Rad Antibodies, Kidlington, UK), TNF-α-BV421 mAb (clone Mab11, BioLegend), IL-2 mAb (clone A150D 3F1 2H2, Invitrogen, Thermo Fisher Scientific) and IL-4 BV605 mAb (clone MP4-25D2, BioLegend) followed by staining with anti-mouse IgG2a-PE-Cy7 (clone RMG2a-62, BioLegend). Cells were analysed using a BD LSRFortessa flow cytometer and FlowJo X software. Total SARS-CoV-2 S specific cytokine positive responses are presented after subtraction of the background response detected in the media stimulated control PBMC sample of each pig, prior to summing together the frequency of S-peptide pools 1-3 specific cells.

### Data Analysis

GraphPad Prism 8.1.2 (GraphPad Software, San Diego, USA) was used for graphical and statistical analysis of data sets. ANOVA or a mixed-effects model were conducted to compare responses over time and between vaccine groups at different time points post-vaccination as detailed in the Results. Neutralising antibody titre data were log transformed before analysis. Neutralising antibody titre data generated by the VNT and pVNT assays were compared using Spearman nonparametric correlation. *p*-values < 0.05 were considered statistically significant.

## Acknowledgements

This study was supported by Engineering and Physical Sciences Research Council (EPSRC) award EP/R013756/1 (VaxHub), UKRI Biotechnology and Biological Sciences Research Council (BBSRC) awards BBS/E/I/00002035, BBS/E/I/00007031 and BBS/E/I/00007039 and the Bill and Melinda Gates Foundation supported ‘Pirbright Livestock Antibody Hub’ (Grant No. OPP1215550). Development of SARS-CoV-2 reagents was partially supported by the NIAID Centers of Excellence for Influenza Research and Surveillance (CEIRS) contract HHSN272201400008C and EPSRC Grant No. EP/S025243/1 to the Rosalind Franklin Institute. We thank V. Clark, H. Gray, and R. Snaith for animal husbandry and the Jenner Institute Vector Core Facility for assistance, and The Pirbright Institute Animal Services Team for animal care and provision of samples.

## Author contributions

Conceptualisation, S.P.G., R.W., J.A.H., E.T., B.C., S.C.G., T.J.T., T.L.; Methodology, S.P.G., R.K.M., A.J.S., S.B.R., J.C.E., J.H., V.M., G.W., C.B., A.L., I.D., H.S., N.T., C.C., D.B.; Investigation, S.P.G., R.K.M., A.J.S., S.B.R., M.U., J.C.E., J.W.P.H., V.M., A.L., G.W., C.B., I.D., H.S., D.B., S.B., P.S.L., P.H., N.T., C.Ch., C.G., D.P., K.M., D.W., H.S., E.A.; Resources, V.M., C.C., J.N., A.S.A., A.B., S.C., J.H., R.J.O., M.C.; Writing—Original Draft Preparation, S.P.G., S.C.G. and T.L.; Writing—Review and Editing, All authors; Project Administration, S.P.G., R.W., S.C.G., T.J.T., T.L.; Funding Acquisition, J.A.H., B.C., S.C.G. T.J.T., and T.L.

## Materials & Correspondence

Correspondence and material requests to Professor Simon P. Graham and Professor Teresa Lambe.

## References

1 Zhu, F.-C. et al. Safety, tolerability, and immunogenicity of a recombinant adenovirus type-5 vectored COVID-19 vaccine: a dose-escalation, open-label, non-randomised, first-in-human trial. The Lancet, doi: https://doi.org/10.1016/S0140-6736(20)31208-3 (2020).

2 van Doremalen, N. et al. A single-dose ChAdOx1-vectored vaccine provides complete protection against Nipah Bangladesh and Malaysia in Syrian golden hamsters. PLoS Negl Trop Dis 13, e0007462, doi: 10.1371/journal.pntd.0007462 (2019).

3 Folegatti, P. M. et al. Safety and immunogenicity of a candidate Middle East respiratory syndrome coronavirus viral-vectored vaccine: a dose-escalation, open-label, non-randomised, uncontrolled, phase 1 trial. The Lancet. Infectious diseases, doi: 10.1016/s1473-3099(20)30160-2 (2020).

4 Munster, V. J. et al. Protective efficacy of a novel simian adenovirus vaccine against lethal MERS-CoV challenge in a transgenic human DPP4 mouse model. NPJ Vaccines 2, 28–28, doi: 10.1038/s41541-017-0029-1 (2017).

5 van Doremalen, N. et al. A single dose of ChAdOx1 MERS provides broad protective immunity against a variety of MERS-CoV strains. bioRxiv, 2020.2004.2013.036293, doi: 10.1101/2020.04.13.036293 (2020).

6 van Doremalen, N. et al. ChAdOx1 nCoV-19 vaccination prevents SARS-CoV-2 pneumonia in rhesus macaques. bioRxiv, 2020.2005.2013.093195, doi: 10.1101/2020.05.13.093195 (2020).

7 van Doremalen, N. et al. A single dose of ChAdOx1 MERS provides protective immunity in rhesus macaques. Science Advances, eaba8399, doi: 10.1126/sciadv.aba8399 (2020).

8 Suleman, M. et al. Antigen encoded by vaccine vectors derived from human adenovirus serotype 5 is preferentially presented to CD8+ T lymphocytes by the CD8α+ dendritic cell subset. Vaccine 29, 5892–5903, doi: 10.1016/j.vaccine.2011.06.071 (2011).

9 Zhang, Y. et al. Immunization with an adenovirus-vectored TB vaccine containing Ag85A-Mtb32 effectively alleviates allergic asthma. J Mol Med (Berl) 96, 249–263, doi: 10.1007/s00109-017-1614-5 (2018).

10 Meurens, F., Summerfield, A., Nauwynck, H., Saif, L. & Gerdts, V. The pig: a model for human infectious diseases. Trends in microbiology 20, 50–57, doi: 10.1016/j.tim.2011.11.002 (2012).

11 Gerdts, V. et al. Large Animal Models for Vaccine Development and Testing. ILAR Journal 56, 53–62, doi: 10.1093/ilar/ilv009 (2015).

12 Rivera-Hernandez, T. et al. The contribution of non-human primate models to the development of human vaccines. Discov Med 18, 313–322 (2014).

13 Holzer, B. et al. Comparison of Heterosubtypic Protection in Ferrets and Pigs Induced by a Single-Cycle Influenza Vaccine. Journal of immunology (Baltimore, Md. : 1950) 200, 4068–4077, doi: 10.4049/jimmunol.1800142 (2018).

14 Holzer, B. et al. Immunogenicity and Protective Efficacy of Seasonal Human Live Attenuated Cold-Adapted Influenza Virus Vaccine in Pigs. Front Immunol 10, 2625, doi: 10.3389/fimmu.2019.02625 (2019).

15 Morgan, S. B. et al. Aerosol Delivery of a Candidate Universal Influenza Vaccine Reduces Viral Load in Pigs Challenged with Pandemic H1N1 Virus. Journal of immunology (Baltimore, Md. : 1950) 196, 5014–5023, doi: 10.4049/jimmunol.1502632 (2016).

16 McLean, R. K. & Graham, S. P. Vaccine Development for Nipah Virus Infection in Pigs. Front Vet Sci 6, 16, doi: 10.3389/fvets.2019.00016 (2019).

17 Pedrera, M. et al. Bovine Herpesvirus-4-Vectored Delivery of Nipah Virus Glycoproteins Enhances T Cell Immunogenicity in Pigs. Vaccines (Basel) 8, doi: 10.3390/vaccines8010115 (2020).

18 Grifoni, A. et al. Targets of T Cell Responses to SARS-CoV-2 Coronavirus in Humans with COVID-19 Disease and Unexposed Individuals. Cell, doi: 10.1016/j.cell.2020.05.015 (2020).

19 Zheng, H. Y. et al. Elevated exhaustion levels and reduced functional diversity of T cells in peripheral blood may predict severe progression in COVID-19 patients. Cell Mol Immunol 17, 541–543, doi: 10.1038/s41423-020-0401-3 (2020).

20 Chen, G. et al. Clinical and immunological features of severe and moderate coronavirus disease 2019. J Clin Invest 130, 2620–2629, doi: 10.1172/jci137244 (2020).

21 Xiong, Y. et al. Transcriptomic characteristics of bronchoalveolar lavage fluid and peripheral blood mononuclear cells in COVID-19 patients. Emerg Microbes Infect 9, 761–770, doi: 10.1080/22221751.2020.1747363 (2020).

22 Braun, J. et al. Presence of SARS-CoV-2 reactive T cells in COVID-19 patients and healthy donors. medRxiv, 2020.2004.2017.20061440, doi: 10.1101/2020.04.17.20061440 (2020).

23 Chandrashekar, A. et al. SARS-CoV-2 infection protects against rechallenge in rhesus macaques. Science, doi: 10.1126/science.abc4776 (2020).

24 Yu, J. et al. DNA vaccine protection against SARS-CoV-2 in rhesus macaques. Science, eabc6284, doi: 10.1126/science.abc6284 (2020).

25 Asthagiri Arunkumar, G. et al. Vaccination with viral vectors expressing NP, M1 and chimeric hemagglutinin induces broad protection against influenza virus challenge in mice. Vaccine 37, 5567–5577, doi: https://doi.org/10.1016/j.vaccine.2019.07.095 (2019).

26 Amanat, F. et al. A serological assay to detect SARS-CoV-2 seroconversion in humans. Nature Medicine, doi: 10.1038/s41591-020-0913-5 (2020).

